# A high-quality genome assembly from short and long reads for the non-biting midge *Chironomus riparius* (Diptera)

**DOI:** 10.1101/659433

**Authors:** Hanno Schmidt, Ann-Marie Waldvogel, Sören Lukas Hellmann, Barbara Feldmeyer, Thomas Hankeln, Markus Pfenninger

## Abstract

**Background:** *Chironomus riparius* is of great importance as a study species in various fields like ecotoxicology, molecular genetics, developmental biology and ecology. However, only a fragmented draft genome exists to date, hindering the recent rush of population genomic studies in this species.

**Findings:** Making use of 50 NGS datasets, we present a hybrid genome assembly from short and long sequence reads that make *C. riparius’* genome one of the most contiguous Dipteran genomes published, the first complete mitochondrial genome of the species and the respective recombination rate as one of the first insect recombination rates at all.

**Conclusions:** The genome and associated resources will be highly valuable to the broad community working with dipterans in general and chironomids in detail. The estimated recombination rate will help evolutionary biologist gain a better understanding of commonalities and differences of genomic patterns in insects.

## Introduction

Non-biting midges (Chironomidae) are dipterans in close phylogenetic relationship to model organisms like *Drosophila* fruit flies and *Anopheles* mosquitoes. Especially the species *Chironomus riparius* (synonym *C. thummi* or *C. thummi thummi*) is important in ecotoxicological [1, 2], molecular genetic [3, 4], developmental [5] and ecological [6–8] research. Recently, *C. riparius* has also emerged as a promising organism for transcriptomic [9–11] and genomic studies [12–14]. Although important population genomic parameters are already available for *C. riparius* (e.g. the mutation rate *μ*; [15]), analyses still rely on a fragmented Illumina-only genome assembly [12]. Here we present a high-quality hybrid genome assembly from short and long reads, along with an estimate for the species-specific recombination rate, the first complete mitochondrial genome for this species and a reference transcriptome based on several life stages. This is an important step forward to enable more complex genomic studies on *C. riparius* and hence understand variability in dipteran genome evolution patterns.

## Assembly strategy

The assembly of the Illumina-PacBio-hybrid genome consisted of five major steps: (1) De Bruijn graph assembly of the Illumina reads, (2) hybrid assembly of Illumina contigs and raw PacBio reads, (3) error correction of hybrid contigs by mapping of Illumina data, (4) scaffolding of the contigs using mate-pair reads, (5) closing the remaining gaps with corrected PacBio and Illumina paired end sequences.

## Samples and PacBio sequencing

Long reads (**Fehler! Verweisquelle konnte nicht gefunden werden.**, dataset 01) were sequenced from 52 female imagos that originated from one egg clutch of a strict inbred line (described in [15]) of the same *C. riparius* laboratory culture that has been used for previous draft genome sequencing [12]. DNA was isolated with the QIAGEN Gentra^®^ Puregene^®^ Tissue Kit according to manufacturer’s instructions and sequenced on six SMRT Cells on a Pacific Biosciences RS II machine.

The 1,155,855 PacBio reads had an average length of 4,751 bp, and the longest read was 48,745 bp.

## Genome assembly

Illumina data (**Fehler! Verweisquelle konnte nicht gefunden werden.**, datasets 02-06) was sequenced from approximately 50 larvae from a long-standing laboratory culture [12]. Quality processing of the reads was done with Trimmomatic v0.32 [16] and FastQC v0.11.3 [17]. Additionally, we filtered out mitochondrial reads using BBDuk from the tool package BBMap v35.85 [18] with k=41 and hdist=2.

We assembled the quality processed Illumina reads using the De Bruijn graph assembler Platanus v1.2.4 [19] with kmer-sizes between k=32 and k=84 and s=6. These contigs plus the raw PacBio reads were then used as input for the program DBG2OLC v1.0 [20] and assembled with recommended settings (k=17, KmerCovTh=2, MinOverlap=20, AdaptiveTh=0.002). The assembly was screened for *Wolbachia* sequences and five contigs were removed, thereby getting rid of all *Wolbachia* contaminations. Since the raw PacBio reads were used for assembly to achieve highest contiguity of contig sequences, we subsequently used proovread v2.13.12 [21] to correct the DBG2OLC contig sequences iteratively. In the first pass, we used the Platanus contig sequences described above, and in a second pass the additional Illumina reads (100x coverage; **Fehler! Verweisquelle konnte nicht gefunden werden.**, datasets 07-11). The Illumina data for error correction was sequenced from progeny of the same one egg clutch as the larvae for the PacBio sequences to allow for highest sequence conformity and thus correction confidence. Since hybrid assembly is a highly complex procedure and we did not want to miss any information, we screened Illumina-only and PacBio-only assemblies for additional sequence information lacking in the DBG2OLC contigs. The Platanus-derived contigs described above were compared to the DBG2OLC contigs by BLASTN searches (Blast v2.3.0+ [22]) with perc_identity=80. The PacBio reads were assembled with Canu v1.0 [23] using default settings and the output contigs (Supplementary Table S2) used for BLASTN searches as described for the Platanus contigs. All contigs from both approaches that did not match DBG2OLC contigs with at least 80% were then added to the DBG2OLC assembly. These sequences were then scaffolded using SSPACE v3.0 [24] with x=0, n=25 and mate-pair libraries with 3 and 5.5 KB insert size (deviation 0.8). Scaffold gaps were addressed with an iterative gap closure process. First, we corrected PacBio raw reads with Illumina reads by proovread applying default settings, and then used them to close gaps with PBJelly v15.2.20 [25]. Afterwards, datasets 02-06 (**Fehler! Verweisquelle konnte nicht gefunden werden.**) were used as input to five iterative rounds of GapFiller v1.10 [26].

The final hybrid assembly consisted of 752 scaffold sequences with a total length of 178,167,951 bp (**Fehler! Verweisquelle konnte nicht gefunden werden.**). The total assembly length fits the published genome size of ~200 Mb estimated by flow cytometry [27], given that regions of low sequence complexity (e.g. highly repetitive parts of centromeres and telomeres) are likely not to be resolved and thus missing. In light of the many tandem-repetitive element clusters interspersed in the genome of *C. riparius* [28, 29] it is therefore reasonable to assume that scaffold ends represent borders to internally repetitive heterochromatic regions in most cases. The N50 of 539,778 bp of the current genome draft is almost twice as high as for a previous version (Table 1). Gap filling drastically reduced the final unresolved base content (N’s) of the assembly down to 0.08%, with the PacBio reads being especially helpful (Figure 1). On average 96.6% of the Illumina sequence reads could be mapped back to the draft genome (Supplementary Table S3), corroborating our assumption that only highly repetitive areas are underrepresented in the genome draft.

**Table 1.** Relative improvement of the assembly for *C. riparius*. Shown are the improvements in quality by combining short and long reads in comparison to the previous Illumina-only assembly. Values are based on nuclear genome only.

|  | Illumina-only draft<br>genome published in [12] | hybrid-assembly from<br>the present study | change |
| --- | --- | --- | --- |
| number of sequences | 5,292 | 752 | ≈ 1/7 |
| total sequence length (bp) | 180,652,019 | 178,167,951 | ≈ equal |
| average sequence length (bp) | 34,136 | 236,926 | ≈ x7 |
| longest sequence (bp) | 2,056,324 | 2,626,431 | ≈ +25% |
| N50 | 272,065 | 539,778 | ≈ x2 |
| N content (%) | 15.96 | 0.08 | ≈ 1/200 |
| BUSCOs found | 92.8 | 93.7 | ≈ +1% |

**Figure 1.**
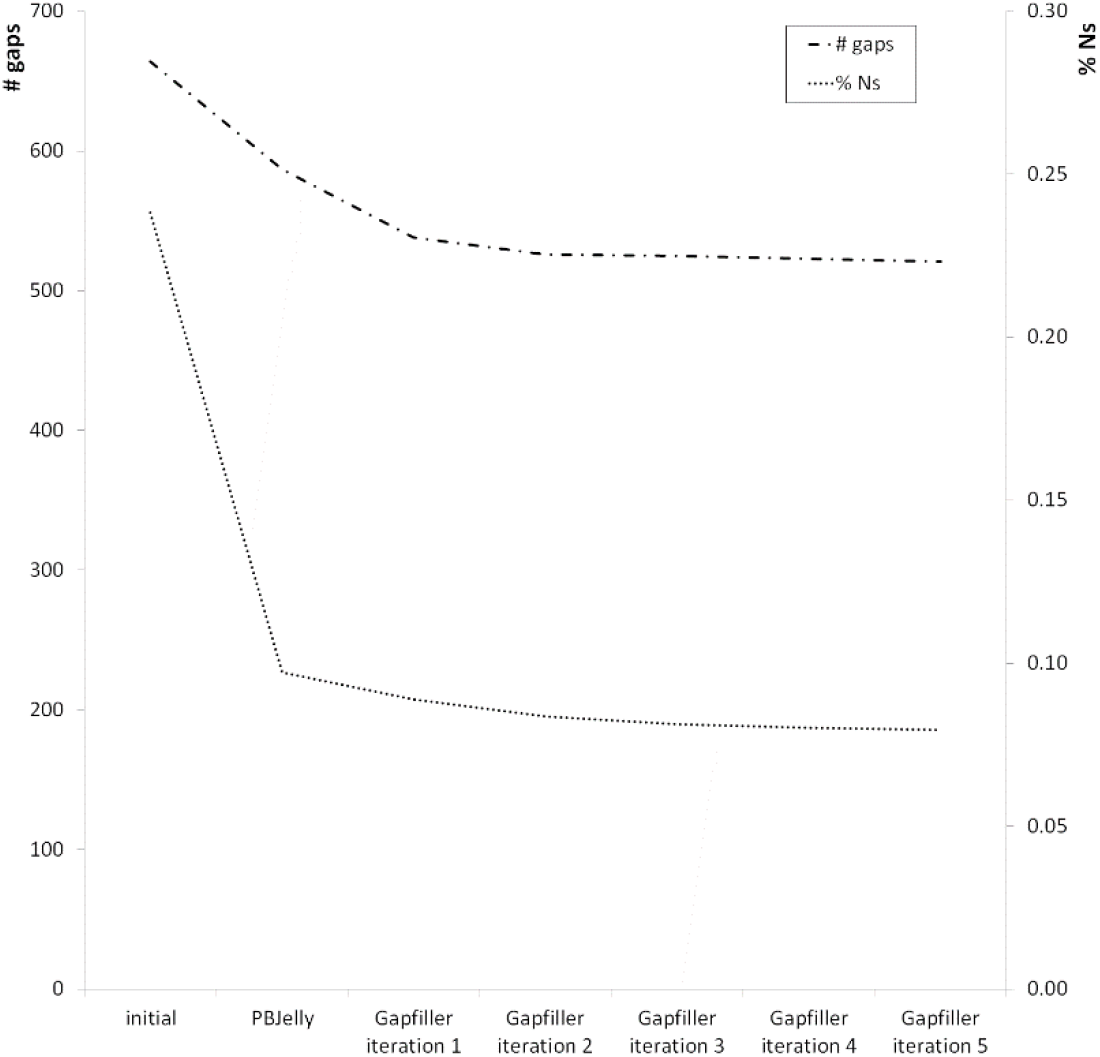
Effect of gap filling procedures. Gap filling with Illumina paired end reads (Gapfiller) and corrected PacBio reads (PBJelly). Shown is the decrease of number of gaps and percentage of N’s in the scaffolds during the iterative gap filling process.

## Estimation of the recombination rate

Using ancestral linkage disequilibrium (LD) based methods for the estimation of recombination rates heavily profits from a genetic map provided at the stage of phasing the SNP data. Since there is no such a resource for *C. riparius*, a constant rate was used. Although this is the default of the phasing algorithm applied, this may introduce a bias into the estimation of *rho* based on this data.

*Rho* values were estimated from 20 field isolates (Supplementary Table S1, datasets 31-50) by applying a *reversible jump Markov Chain Monte Carlo mechanism* (rjMCMC) implemented in the program LDhelmet v1.7 [30] individually for each scaffold. LDhelmet is a derivative of LDhat [31], especially modified to fit genomic characteristics that differ from hominids to *Drosophila* (for example higher SNP density). Since we anticipate similar patterns in *Chironomus*, we chose LDhelmet and mainly followed the parameter recommendations of the authors. The ultimate LDhelmet analysis with the *rjmcmc* command was run for each scaffold with a block penalty of 50.0 (as recommended; parameter of negligible influence on results [32]) and a window size of 50 SNPs (as in the data preparation). We used a burn-in of 1,000,000 iterations and subsequently ran the Markov chain for 10,000,000 iterations (see Supplementary Methods for details).

Mean *ρ* (always given per base pair within a 50 kb windows) ranged between 0.04 and 0.07. *C. riparius ρ* values therefore lie within the range of those estimated for *Drosophila melanogaster* (0.01 to 0.11; [30]).

Recombination should be less frequent across sex-determining regions, because reciprocal exchange of chromosomal parts is only possible in the germ line of female individuals [33] [but see 34]. *C. riparius* has a sex-determination system with heterogametic males, bearing a sex-determining region (SDR) on chromosome 3 that is being interpreted as an emerging sex chromosome [35, 36]. We identified the SDR by BLASTN searches with the sequence of the single copy gene CpY (NCBI accession number X82317.1) as query. Indeed, the scaffold containing the SDR showed a large region with lowered recombination rates around CpY. When extracting the last 600,000 bp from scaffold 549 (i.e. the potential SDR), we see a mean recombination rate of 0.014 for the SDR compared to values between 0.024 and 0.07 for the four chromosomes (Figure 2). This seems reasonable since recombination should be roughly halved in the SDR. The remaining parts of chromosome 3 without the identified part of the SDR still have relatively low recombination rates (Figure 2), potentially due to further fragments of the SDR on other scaffolds or effects of the SDR on the genomic surrounding. Statistical analysis with Kruskal-Wallis tests as described above confirmed that recombination rates of chromosome 3 were significantly different to those of all other chromosomes (p value < 0.001).

**Figure 2.**
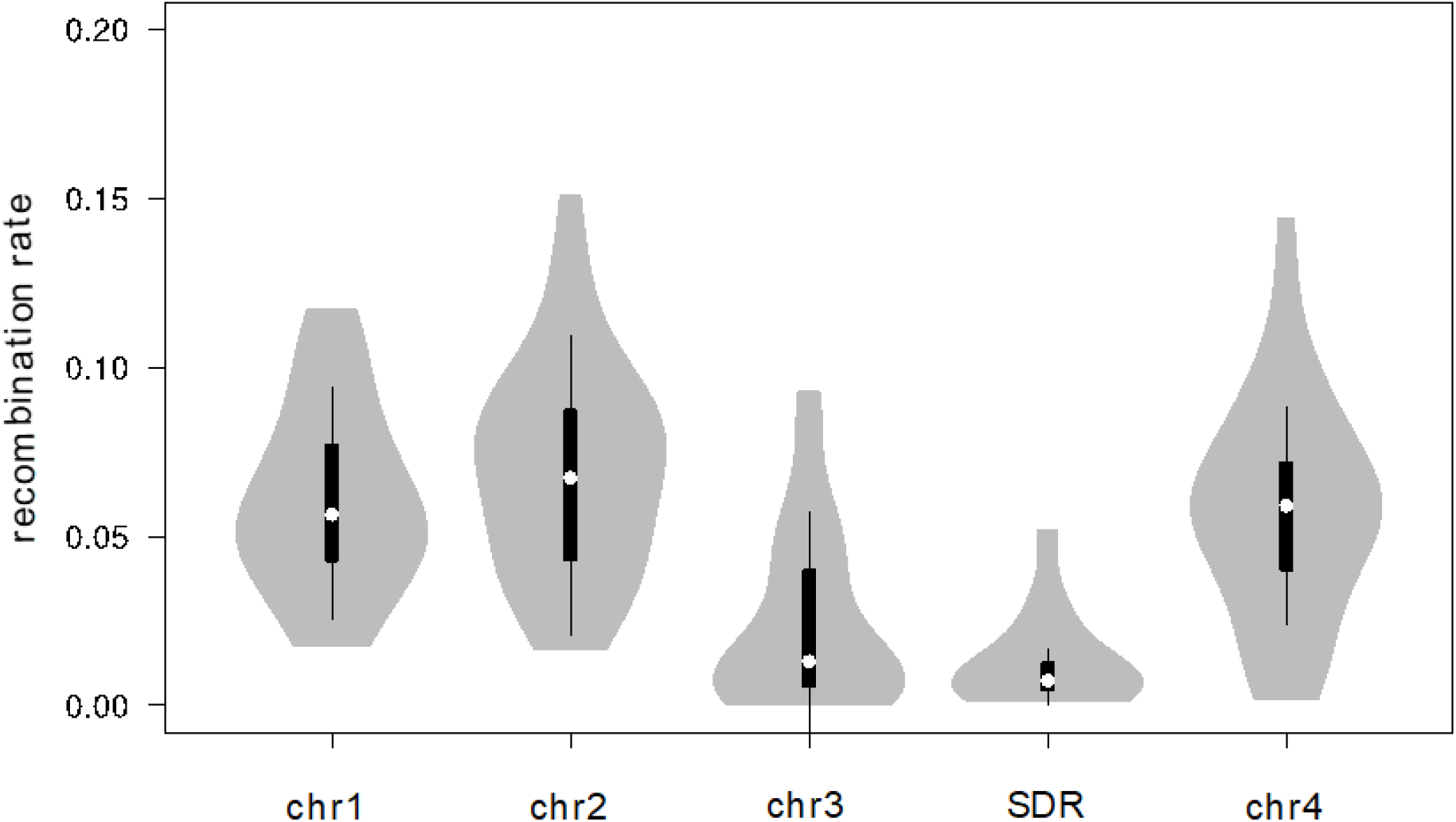
*C. riparius* chromosome-specific recombination rates. Recombination rates from all individuals across populations were pooled. Chr = Chromosome. Chromosome 3 is represented without the identified part of the sex determining region, which is displayed separately (SDR). White circles show medians, box limits indicate the 25th and 75th percentiles, whiskers extend 1.5 times the interquartile range from the 25th and 75th percentiles, polygons represent density estimates of data and extend to extreme values. Kruskal-Wallis test (nonparametric for data without normal distribution) with Dunn’s multiple comparison post-test (GraphPad Prism v5) revealed a significant difference (p < 0.001) between chromosome 3 as well as the SDR relative to all other chromosomes.

## Genome annotation

For the annotation of the gene set of the genome a reference transcriptome was assembled from 19 cDNA sequence data sets (**Fehler! Verweisquelle konnte nicht gefunden werden.**, datasets 12-30). We used Illumina and 454 Roche sequence reads from embryos, larvae and adults (both sexes; treated and untreated; see Supplementary Table S1 and references therein for details) to guarantee for as many expressed genes as possible in order to optimise gene annotation.

Firstly, data sets were pre-processed using fastqc [17] and BBDuk from the BBMap package v35.85 [18]. Thereby, sequence adapters were trimmed using k=23, mink=11, hdist=1, tbo and tpe options. 3’ bases with phred quality below 20 were trimmed and reads with average phred quality below 20 discarded. Assembly of the cleaned reads was then performed in two separate steps for Illumina and 454 data with Trinity v2.3.2 [37] using uneven k-mer sizes from 25 to 31. The best assembly was identified to be with k=25 using assembly metrics like N50 and a search for core orthologous genes with BUSCO v1.2b [38] and used further on. The resulting assemblies for Illumina and 454 Roche data, respectively, were then merged and duplicate contigs removed using dedupe from the BBmap package with mid=90. The resulting final transcriptome assembly was then used for gene annotation.

The whole annotation process was performed with the MAKER2 v2.31.8 [39, 40] pipeline and affiliated programs. Before running MAKER2, all data available for annotation of the genome has to be prepared accordingly and handed to MAKER2 in form of input files. To ensure discovery of most repeat sequences in the draft genome, we extended the custom repeat library from [12] with repeat sequences extracted manually from the draft genome presented in this study.

The draft genome, the reference transcriptome described above and the GFF file from the BUSCO run were uploaded to the Augustus v3.2.3 training [41] at the University of Greifswald webserver (http://bioinf.uni-greifswald.de/webaugustus/training/create; accessed on 2017-02-24). This output then served as input for the first round of MAKER2. MAKER2 can work much more accurate when provided with genome-specific gene models at the beginning. Therefore we ran CEGMA v2.5 [42] on the draft genome and converted the output to a hidden Markov model using scripts from the SNAP gene finder v2006-07-28 [43]. Additionally, we created another hidden Markov model by running GeneMark v4.32 [44] with min_contig set to 20,000 on the draft genome.

The first round of MAKER2 annotation was then run using MPICH2 v3.2 (https://www.mpich.org/) parallelisation with the described transcriptome, SNAP, GeneMark and Augustus models, our custom repeat library plus the SwissProt database (as at 13.1.2016) as input. The MAKER2 pipeline was run with the programs Augustus v3.2.1, BLAST v2.2.28+ [22], Repeatmasker v4.0.6 [45], SNAP v2006-07-28 [43], GeneMark v4.32 [44] and Exonerate v2.2.0 [46]. We applied default parameters with only max_dna_len set to 500,000 to prevent loss of gene parts from genes with larger introns, min_protein set to 10 to receive as much potential protein sequences as possible and fix_nucleotides set as flag to allow for non-ACGT-characters in the genome file. The gff files of MAKER2’s output were merged to a single file using gff3_merge from the MAKER2 distribution. Afterwards this gff file was converted to a hidden Markov model using SNAP scripts as described above for CEGMA with only the cegma2zff script being replaced by maker2zff. The information from the first MAKER2 run was also used for retraining the gene model in Augustus. The genome.ann file was first retransformed to gff and modified to match the gff format and then fed into the autoAug.pl script from Augustus v3.2.1 for retraining the EST-based gene models. The second round of the MAKER2 pipeline was then started with the same settings and input files as described above but with the updated Augustus and SNAP gene models and the parameters min_protein set to 30 and alt_splice on. Afterwards, a third round of the MAKER2 pipeline was run exactly as the second one, including another re-training of the Augustus gene model with autoAug.pl and again updating the SNAP gene model. From the resulting output of MAKER2’s third round we merged all gff files with gff3_merge as described above and renamed the included gene tracks to ensure easier handling. To allow for assigning putative gene functions to the annotated gene tracks, we gathered all predicted proteins from the MAKER2 output folder, performed BLASTP searches against the SwissProt database (as at 13.1.2016) and then added the best BLAST hits to the accordant gene tracks.

13,449 protein-coding genes were annotated across the nuclear genome (Table 2). Applying the algorithm BUSCO v1.2b [38], we found orthologous sequences to 93.7% of arthropod core genes (Table 1), we can assume the genome to be almost complete also in terms of gene space. 101,693 regions up to 30 kb in length were annotated as repetitive sequences (9.14%). Compared to the Illumina-only genome draft [12], the inclusion of long sequence reads has significantly increased detection of repeats by 41%. Given the heavy load of repetitive sequences in the *C. riparius* genome [47] however, this value most likely still represents an underestimate due to unresolved large heterochromatic regions.

**Table 2.** Annotation of the nuclear genome. Content of protein-coding genes in the *C. riparius* genome compared to *C. tentans* and *D. melanogaster*.

|  | gene count | average number of exons per gene | average exon length | protein coding part of the genome |
| --- | --- | --- | --- | --- |
| *C. riparius* | 13,449 | 5.1 | 378 bp | 14.6% |
| *C. tentans* | 15,120 | 3.8 | 312 bp | 9% |
| *D. melanogaster* | 13,907 | 5.5 | 538 bp | 18.3% |

## Assembly and annotation of the mitochondrial genome

The reconstruction of the mitochondrial genome sequence was performed using the program MITObim v1.8 [48] on the large paired end dataset 03. MITObim applies a baiting and iterative mapping approach to short read data. Using a mitochondrial reference sequence (here an unpublished, partial Sanger sequence of *C. riparius’* mitochondrial genome), the program performs a mapping to gather all reads belonging to the mitochondrial genome and assembles them with MIRA v4.0.2 [49] in the first round. Then it uses the produced sequence to again fish for mitochondrial reads for assembly. This is repeated until the number of mapped reads becomes stationary. MITObim was run four times with the reference sequences of *C. riparius* being modified in length to allow for different starting points of the procedure. All four output sequences were then aligned in MEGA v7.0.7 [50] and manually integrated into a consensus sequence. This consensus sequence was annotated by MITOS WebServer [51] using the genetic code 05 - invertebrate. The whole sequence and all annotations were finally checked and, where necessary, corrected manually.

The mitochondrial genome’s length is 15,467 bp, which is in line with other dipteran values. All 37 genes of the mitochondrial genome could be annotated and defined exactly (Figure 3, Table 3).

**Figure 3.**
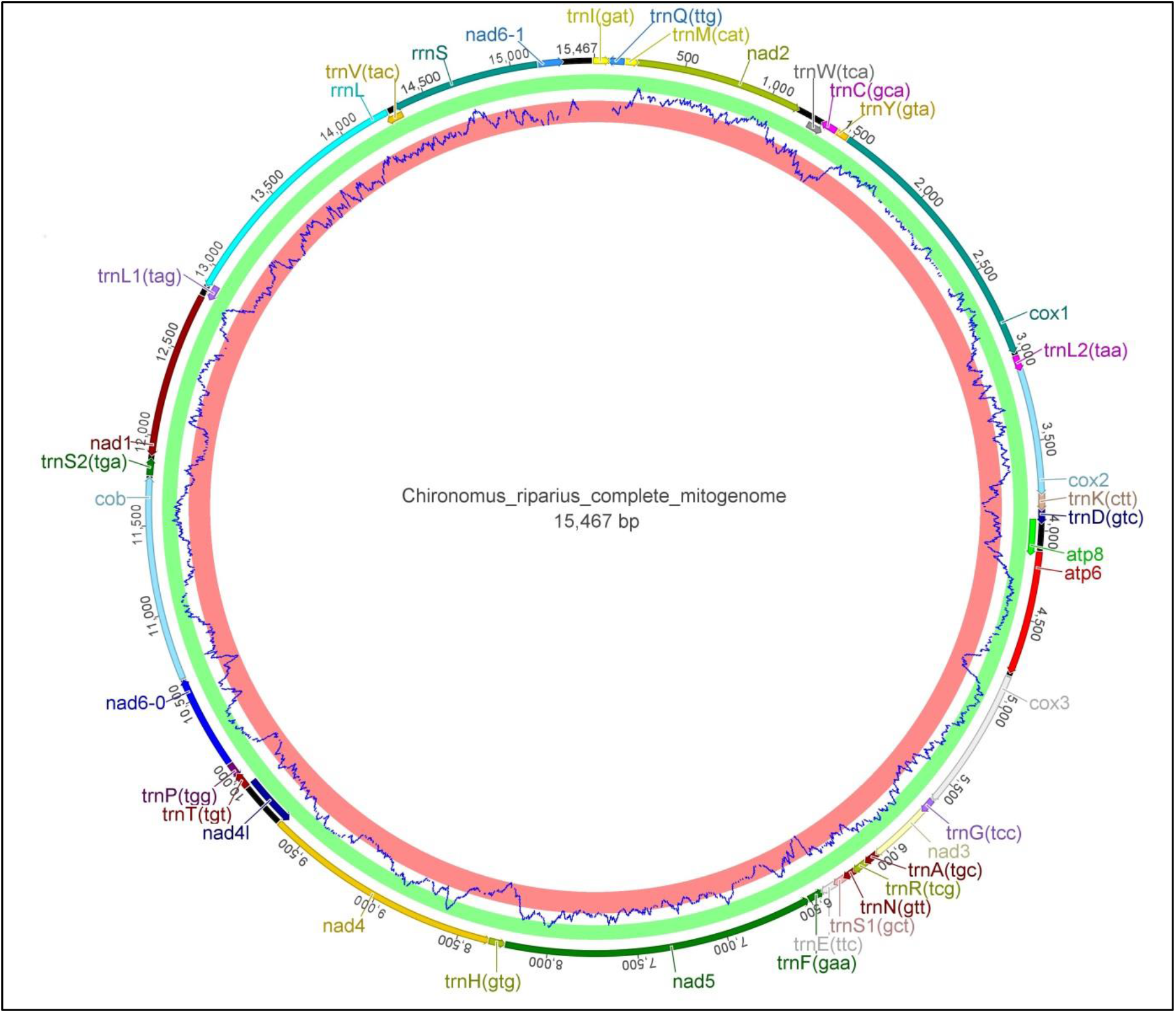
Mitochondrial genome. The circular genome consists of 15,467 bp. Prediction of protein-coding sequence parts from the EMBOSS tool tcode (graph at the inner edge of the genome) mainly is consistent with the annotation from MITOS.

**Table 3.** Annotation of the mitochondrial genome. The annotation was generated by MITOS and manually curated afterwards.

| Gene name | Start | Stop | Strand |
| --- | --- | --- | --- |
| trnI(gat) | 1 | 68 | + |
| trnQ(ttg) | 82 | 150 | - |
| trnM(cat) | 159 | 227 | + |
| nad2 | 249 | 1154 | + |
| trnW(tca) | 1256 | 1323 | + |
| trnC(gca) | 1323 | 1391 | - |
| trnY(gta) | 1398 | 1463 | - |
| cox1 | 1474 | 2988 | + |
| trnL2(taa) | 3028 | 3093 | + |
| cox2 | 3105 | 3785 | + |
| trnK(ctt) | 3802 | 3873 | + |
| trnD(gtc) | 3880 | 3948 | + |
| atp8 | 3949 | 4125 | + |
| atp6 | 4125 | 4790 | + |
| cox3 | 4837 | 5616 | + |
| trnG(tcc) | 5639 | 5704 | + |
| nad3 | 5705 | 6049 | + |
| trnA(tgc) | 6060 | 6125 | + |
| trnR(tcg) | 6138 | 6204 | + |
| trnN(gtt) | 6205 | 6272 | + |
| trnS1(gct) | 6273 | 6339 | + |
| trnE(ttc) | 6347 | 6415 | + |
| trnF(gaa) | 6432 | 6498 | - |
| nad5 | 6522 | 8222 | - |
| trnH(gtg) | 8247 | 8313 | - |
| nad4 | 8335 | 9657 | - |
| nad4l | 9654 | 9935 | - |
| trnT(tgt) | 9949 | 10014 | + |
| trnP(tgg) | 10015 | 10081 | - |
| nad6-0 | 10104 | 10613 | + |
| cob | 10642 | 11754 | + |
| trnS2(tga) | 11793 | 11860 | + |
| nad1 | 11902 | 12801 | - |
| trnL1(tag) | 12826 | 12892 | - |
| rrnL | 12888 | 14267 | - |
| trnV(tac) | 14266 | 14337 | - |
| rrnS | 14337 | 15138 | - |
| nad6-1 | 15140 | 15280 | + |

## Author’s contributions

MP, TH and HS conceived the study. HS and SLH assembled and annotated the genome. HS, A-MW and BF estimated recombination rates. All authors interpreted the data. HS drafted the manuscript. All authors contributed to the final version of the manuscript.

## Acknowledgements

We gratefully acknowledge support in LDhelmet usage by Paul Jenkins and Mathias Weber and assistance in genome annotation by Florian Dolze. Parts of this research were conducted using the supercomputer Mogon and/or advisory services offered by Johannes Gutenberg University Mainz (hpc.uni-mainz.de), which is a member of the AHRP and the Gauss Alliance e.V.

