## Supplements for "A high-quality genome assembly from short and long reads for the non-biting midge *Chironomus riparius* (Diptera)"

---

---

### Supplementary Methods: Estimation of the recombination rate

Short read sequence data from twenty unrelated female individuals (datasets 27-46 from Supplementary Table S1, published in [1]) from five populations across Europe was used to generate a set of single nucleotide polymorphisms (SNP) for the analyses. Sampling locations and quality processing of the reads are described in [2].

Reads were aligned to the genome described in this study using bwa mem algorithm from BWA v0.7.13-r1126 [3, 4] with standard parameters. PCR duplicate reads were marked with the MarkDuplicatesWithMateCigar algorithm from Picard tools v2.6.0 (http://broadinstitute.github.io/picard/). Afterwards, realignment around indels was performed with the IndelRealigner algorithm from the Genome Analysis Toolkit v3.6-0-g89b7209 (GATK; [5]), followed by recalibration of bases. For the latter we applied the UnifiedGenotyper algorithm from GATK to identify an initial SNP set, searched for shared SNPs in all resulting VCF files with the isec algorithm from Bcftools v1.3.1 [6] and finally recalibrated bases with the BaseRecalibrator algorithm from GATK.

The final SNPs were called on the recalibrated BAM files again using the *UnifiedGenotyper* algorithm from GATK with *stand_call_conf*=40 and *stand_emit_conf*=15. Phasing of haplotypes was performed scaffold-wise with SHAPEIT2 v2.r837 [7] using only diallelic SNPs. We converted the output from SHAPEIT2 to VCF files and extracted input files for LDhelmet with VCFtools (http://vcftools.sourceforge.net) and the *--ldhelmet* option.

Recombination rates were estimated by applying a *reversible jump Markov Chain Monte Carlo mechanism* (rjMCMC) implemented in the program LDhelmet v1.7 [8] individually for each scaffold. LDhelmet is a derivative of LDhat [9], especially modified to fit genomic characteristics that differ from hominids to *Drosophila* (for example higher SNP density). Since we anticipate similar patterns in *Chironomus*, we chose LDhelmet and mainly followed the parameter recommendations of the authors.

Using the information from the SNPS and POS files from the VCFtools' output we generated FASTA files for every population and every scaffold individually, ending up with 3,220 input files (5 populations * 644 scaffolds), containing 25,760 haplotype blocks (3,220 files * 4 individuals * 2 haplotypes), representing 99 % of the genome assembly. Data preparation for the main run in LDhelmet included creation of haplotype configuration files with the *find_confs* command on a window size of 50 SNPs as recommended by the authors, as well as likelihood lookup tables and Padé coefficients. For these calculations we used population-scaled mutation rates *theta* ($\theta = 4N_e\mu$) of $\theta_{MF} = 0.01236$, $\theta_{MG} = 0.01134$, $\theta_{NMF} = 0.01446$, $\theta_{SI} = 0.01029$ and $\theta_{SS} = 0.01021$, that were derived from Pool-Seq data of the same populations [1] and calculations with PoPoolation v1.2.2 [10]. Default values were applied to all other parameters.

The ultimate LDhelmet analysis with the *rjmcmc* command was run for each scaffold with a block penalty of 50.0 (as recommended; parameter of negligible influence on results [11]) and a window size of 50 SNPs (as in the data preparation). We used a burn-in of 1,000,000 iterations and subsequently ran the Markov chain for 10,000,000 iterations (in addition to the burn-in). Results were extracted from the binary output via the *post_to_text* command of LDhelmet.

### Supplementary Tables

**Supplementary Table S1 – Sequence data used in the study**

| dataset ID | used in part | sequencing technology | read length (bp) | number of raw reads | accession number | publication |
|---|---|---|---|---|---|---|
| 01 | genome assembly | PacBio RS II | up to 48,745 | 1,155,855 | pending | **this study** |
| 02 | genome assembly | Illumina paired end | 100 | 29,136,088 | SAMEA4560833 | [2] |
| 03 | genome assembly | Illumina paired end | 100 | 167,264,372 | SAMEA4560834 | [2] |
| 04 | genome assembly | Illumina paired end (MiSeq) | 300 | 58,447,092 | SAMEA4560835 | [2] |
| 05 | genome assembly + scaffolding | Illumina mate-pair | 100 (3 kb insert size) | 48,853,530 | SAMEA4560836 | [2] |
| 06 | genome assembly + scaffolding | Illumina mate-pair | 100 (6 kb insert size) | 47,978,148 | SAMEA4560837 | [2] |
| 07 | error correction | Illumina paired end | 150 | 29,694,824 | PRJEB18039 | [12] |
| 08 | error correction | Illumina paired end | 150 | 34,118,424 | PRJEB18039 | [12] |
| 09 | error correction | Illumina paired end | 150 | 35,803,108 | PRJEB18039 | [12] |
| 10 | error correction | Illumina paired end | 150 | 38,534,360 | PRJEB18039 | [12] |
| 11 | error correction | Illumina paired end | 150 | 39,512,980 | PRJEB18039 | [12] |
| 12 | annotation (transcriptome) | 454 Roche | up to 679 | 430,916 | SRR834592 | [13] |
| 13 | annotation (transcriptome) | 454 Roche | up to 914 | 456,584 | SRR834593 | [13] |
| 14 | annotation (transcriptome) | 454 Roche | up to 690 | 1,549,146 | SRR496839 | [14] |
| 15 | annotation (transcriptome) | 454 Roche | up to 606 | 4,041 | SRX022389 | [15] |
| 16 | annotation (transcriptome) | 454 Roche | up to 679 | 138,103 | SRR1049908 | |
| 17 | annotation (transcriptome) | 454 Roche | up to 802 | 266,129 | SRR1049909 | |
| 18 | annotation (transcriptome) | 454 Roche | up to 600 | 189,271 | SRR1049910 | |
| 19 | annotation (transcriptome) | 454 Roche | up to 662 | 235,202 | SRR1049911 | |
| 20 | annotation (transcriptome) | Illumina paired end | 100 | 6,863,480 | SRR1028867 | [16] |
| 21 | annotation (transcriptome) | Illumina paired end | 100 | 10,222,374 | SRR1032319 | [16] |
| 22 | annotation (transcriptome) | Illumina paired end | 100 | 9,684,342 | SRR1032320 | [16] |
| 23 | annotation (transcriptome) | Illumina paired end | 100 | 9,305,342 | SRR1032321 | [16] |
| 24 | annotation (transcriptome) | Illumina paired end | 100 | 10,861,756 | SRR1032322 | [16] |

| dataset ID | used in part | sequencing technology | read length (bp) | number of raw reads | accession number | publication |
|---|---|---|---|---|---|---|
| 25 | annotation (transcriptome) | Illumina paired end | 100 | 10,783,776 | SRR1032323 | [16] |
| 26 | annotation (transcriptome) | Illumina paired end | | | unpublished | 1KITE project |
| 27 | annotation (transcriptome) | Illumina paired end | 100 | 105,429,480 | ERS1472439 | [2] |
| 28 | annotation (transcriptome) | Illumina paired end | 100 | 197,899,590 | ERS1472440 | [2] |
| 29 | annotation (transcriptome) | Illumina paired end | 100 | 34,987,537 | ERS1472441 | [2] |
| 30 | annotation (transcriptome) | Illumina paired end | 100 | 46,557,991 | ERS1472442 | [2] |
| 31 MF1 | recombination rate populations | Illumina paired end | 150 | 36,869,618 | ERR2528543 | [1] |
| 32 MF2 | recombination rate populations | Illumina paired end | 150 | 30,387,616 | ERR2528544 | [1] |
| 33 MF3 | recombination rate populations | Illumina paired end | 150 | 32,170,704 | ERR2528545 | [1] |
| 34 MF4 | recombination rate populations | Illumina paired end | 150 | 32,312,656 | ERR2528546 | [1] |
| 35 MG2 | recombination rate populations | Illumina paired end | 150 | 29,057,938 | ERR2528547 | [1] |
| 36 MG3 | recombination rate populations | Illumina paired end | 150 | 28,788,518 | ERR2528548 | [1] |
| 37 MG4 | recombination rate populations | Illumina paired end | 150 | 27,094,554 | ERR2528549 | [1] |
| 38 MG5 | recombination rate populations | Illumina paired end | 150 | 34,100,258 | ERR2528550 | [1] |
| 39 NMF1 | recombination rate populations | Illumina paired end | 150 | 38,469,060 | ERR2528551 | [1] |
| 40 NMF2 | recombination rate populations | Illumina paired end | 150 | 32,269,454 | ERR2528552 | [1] |
| 41 NMF3 | recombination rate populations | Illumina paired end | 150 | 29,389,106 | ERR2528553 | [1] |
| 42 NMF4 | recombination rate populations | Illumina paired end | 150 | 26,446,596 | ERR2528554 | [1] |
| 43 SI1 | recombination rate populations | Illumina paired end | 150 | 37,946,848 | ERR2528555 | [1] |
| 44 SI2 | recombination rate populations | Illumina paired end | 150 | 42,327,576 | ERR2528556 | [1] |

| dataset ID | used in part | sequencing technology | read length (bp) | number of raw reads | accession number | publication |
|---|---|---|---|---|---|---|
| 45<br>SI3 | recombination rate populations | Illumina paired end | 150 | 32,922,356 | ERR2528557 | [1] |
| 46<br>SI4 | recombination rate populations | Illumina paired end | 150 | 30,872,238 | ERR2528558 | [1] |
| 47<br>SS1 | recombination rate populations | Illumina paired end | 150 | 31,078,600 | ERR2528559 | [1] |
| 48<br>SS2 | recombination rate populations | Illumina paired end | 150 | 39,785,504 | ERR2528560 | [1] |
| 49<br>SS3 | recombination rate populations | Illumina paired end | 150 | 36,018,158 | ERR2528561 | [1] |
| 50<br>SS4 | recombination rate populations | Illumina paired end | 150 | 28,776,202 | ERR2528562 | [1] |

**Supplementary Table S2 – Statistics for the PacBio-only assembly with Canu**

| | |
|---|---|
| number of sequences | 8,488 |
| total sequence length (bp) | 229,089,447 |
| average sequence length (bp) | 26,990 |
| longest sequence (bp) | 1,085,725 |
| N50 | 56,198 |

**Supplementary Table S3 – Back-mapping of reads used to obtain the assembly onto the draft genome**

Only intact pairs after all quality processing steps were mapped.

| dataset ID | mapped read pairs | % mapped read pairs |
|---|---|---|
| 02 | 6,363,845 | 96.9 |
| 03 | 42,770,555 | 98.1 |
| 04 | 25,635,946 | 99.5 |
| 05 | 13,462,849 | 93.3 |
| 06 | 18,452,418 | 92.2 |
